# A curated *C. difficile* strain 630 metabolic network: prediction of essential targets and inhibitors

**DOI:** 10.1101/006932

**Authors:** Mathieu Larocque, Thierry Chénard, Rafael Najmanovich

## Abstract

*Clostridium difficile* is the leading cause of hospital-borne infections occurring when the natural intestinal flora is depleted following antibiotic treatment. We present *i*MLTC804cdf, an extensively curated reconstructed metabolic network for the *C. difficile* pathogenic strain 630. *i*MLTC804cdf contains 804 genes, 705 metabolites and 766 metabolic, 145 exchange and 118 transport reactions. *i*MLTC804cdf is the most complete and accurate metabolic reconstruction of a gram-positive anaerobic bacteria to date. We validate the model with simulated growth assays in different media and carbon sources and use it to predict essential genes. We obtain 88.8% accuracy in the prediction of gene essentiality when compared to experimental data for *B. subtilis* homologs. We predict the existence of 83 essential genes and 68 essential gene pairs, a number of which are unique to *C. difficile* and have non-existing or predicted non-essential human homologs. For 19 of these potential therapeutic targets, we find 72 inhibitors of homologous proteins that could serve as starting points in the development of new antibiotics, including approved drugs with the potential for drug repositioning.

Systems & Synthetic Biology Subject Category: Genome Scale & Integrative Biology

Molecular & Cell Biology Subject Category: Pharmacology & Drug Discovery

## INTRODUCTION

*Clostridium difficile* is an opportunistic, gram-positive anaerobic spore-forming pathogen found in the environment and in the intestinal flora in up to 3% of healthy adults. Toxigenic strains of *C. difficile* are resistant to a wide variety of antibiotics and produce the enterotoxin TcdA and the cytotoxin TcdB. These toxins are responsible for the clinical symptoms of *C. difficile* infection (CDI) (Voth & Ballard, 2005; Rupnik *et al*, 2009). CDI is the leading cause of hospital-borne infections occurring when the natural intestinal flora is depleted following antibiotic treatment. CDI is the major cause of antibiotic-associated diarrhea and is responsible for pseudomembranous colitis, a form of severe intestinal inflammation. For the most part, CDI can still be treated with metronidazole or vancomycin for which resistance levels remain low or the recently approved fidaxomicin. Single or multiple relapses after initial treatment are common and bring about more severe symptoms. A recent clinical study reports a relapse rate of 24% and 13% with vancomycin or fidaxomicin treatment respectively (Louie *et al*, 2011). Between 50% and 80% of recurrences are due to spore-mediated re-infection (Rupnik *et al*, 2009). Unfortunately, patient-to-patient transmission and relapses are difficult to prevent due to the production of *C. difficile* spores that are resistant to antibiotics, heat, radiation and various chemicals.

CDI is directly responsible for an average 4.6 per 1000 patients admitted in hospitals with a 5.7% mortality rate after 30 days directly attributed to CDI (Gravel *et al*, 2009). In the US, over 250,000 cases are registered per year in hospitals alone and many more cases in outpatient settings (DuPont, 2009), costing around USD$4,000 to USD$8,000 per case of primary infection and USD$8,000 to USD$15,000 per relapsing infection (Ghantoji *et al*, 2010) leading to a burden of over USD$500 million (McGlone *et al*, 2012). More important than economic costs, CDI in older patients, those with concurrent debilitating conditions, and severely relapsing or fulminant cases may result in death.

In recent years there has been an increase in the rate of infections as well as the emergence of community-associated, virulent and antibiotic-resistant strains (Kelly & LaMont, 2008; Shah *et al*, 2010). Treatments of CDI that offer alternatives to the use of small-molecules (Rea *et al*, 2013) involving phages (Hargreaves & Clokie, 2014) or intestinal microbial flora transplants (Brown, 2014) are likely to meet resistance from patients. Others involving antibodies (Humphreys & Wilcox, 2014) or vaccines (Leuzzi *et al*, 2014) are still under development.

The complexity inherent to preventing and treating CDI requires the continuous search for new ways to target *C. difficile*. In recent year, the growth of biological databases led to the development of the field of systems biology making it possible to build and analyze genomic-scale reconstructed metabolic networks (Palsson, 2006). There is a large number of highly curated reconstructed metabolic networks for a number of organisms, from *E. coli* (Reed *et al*, 2003) to human (Thiele *et al*, 2013). The prediction of essential genes is often used to detect potential drug targets (Ghosh *et al*, 2013; Xu *et al*, 2011; Harrold *et al*, 2013). Two techniques, Flux Balance Analysis (FBA) (Orth *et al*, 2010) and Synthetic Accessibility (SA) (Wunderlich & Mirny, 2006) are among those available to predict essential genes at a genomic scale through *in silico* gene deletion studies. The comparison of results obtained with either FBA or SA and experimentally determined essential genes shows equivalent levels of accuracy with either technique around 94% for *B. subtilis* (Oh *et al*, 2007), 83% for *S. cerevisiae* and 60-70% for *E. coli* (Wunderlich & Mirny, 2006). The success rates are likely reflecting the quality of the metabolic network reconstructions.

The combination of systems pharmacology and metabolic network analyses can help predict off-target effects of drugs as well as open new opportunities with the repositioning of existing drugs (Xie *et al*, 2011). In the present study we create and validate a highly curated metabolic network reconstruction for the pathogenic *C. difficile* strain 630. We then utilize it to predict essential genes or gene pairs and detect small-molecules, including existing approved drugs that may bind a number of these targets. We also employ systems biology, bioinformatics and structural computational biology methods to detect potential human cross-reactivity targets.

## RESULTS

### Creation of the network

The genome of *C. difficile* strain 630 is composed of a circular chromosome of 4,290,252 bp coding for 3979 open reading frames (ORFs) as well as a plasmid of 7881 bp coding for 11 ORFs (Sebaihia *et al*, 2006). The *C. difficile* strain 630 draft reconstructed metabolic network presented here covers 20.2% of the ORFs present in the chromosomal genome of the bacteria with 804 ORFs. These 804 genes code for proteins catalyzing 911 reactions (766 metabolic and 145 transport reactions). The model also includes an additional 118 exchange reactions. A total of 593 unique metabolites (705 in total not considering extracellular or intracellular state) are involved in the 911 reactions in the network. The coverage of the genome is similar to those of previously published reconstructed metabolic networks such as *B. subtilis* with 20% (Oh *et al*, 2007) and higher than *C. acetobutylicum* with 12.6% (Senger & Papoutsakis, 2008). Most reactions have at least one gene association (78.07%). Reactions without any gene association were added based on the existence of evidence from the literature such as in the case of Stickland reactions (Barker, 1981; Jackson *et al*, 2006; Nisman, 1954; Mead, 1971), presence in databases such as xanthine amido hydrase or to fill functional gaps to obtain a functional network as in the case of putative transporters for end-products of fermentation.

The final version of the network is available in 3 different formats: 1. An excel file that shows on different spreadsheets the reactions, metabolites, genes, and compartments that comprise the network and the definitions of the network based on the standard described in the RAVEN toolbox (Agren *et al*, 2013). This file is meant to be easily readable by humans; 2. A tab-separated format of the network amenable to analysis in the R Statistical Computing environment (www.r-project.org) using Sybil (Gelius-Dietrich *et al*, 2013); and lastly, 3. A SBML Level 2 formatted network (Hucka *et al*, 2003) that can be used with tools such as Matlab or other SBML compliant software. While a naming convention has been suggested recently for metabolic reconstructions (Reed *et al*, 2003), we feel that a naming convention that does not allude to the name of the organism is insufficient. Therefore, in the present work, the *C. difficile* strain 630 metabolic network reconstruction is called *i*MLTC804cdf as per the suggested convention with the added cdf suffix denoting the KEGG (Kanehisa *et al*, 2014) three-letter organism ID representing *C. difficile* strain 630. The SBML version of the model has been deposited to the BioModels database (Li *et al*, 2010) and assigned the identifier MODEL1407050000.

### Validation

Four types of growth media were simulated *in silico* via modulation of the exchange reactions for the import of metabolites present in the simulated growth media. All four tested media (Table I) produced biomass based on FBA and SA analysis (Wunderlich & Mirny, 2006). For SA, 4 proteins (oxidized ferredoxin, oxidized thioredoxin, acyl and sulfur carrier proteins) that cannot be produced due to the absence of protein biosynthesis reactions in the network but at equilibrium in FBA had to be supplemented to the media to make all reactions possible in FBA also accessible in SA. ATP and nicotinate were supplemented to the minimal medium to allow biomass production in SA while ATP was added to the complex medium.

**Table I:** Definition of the different media used in this study

| Medium name | Components<sup>e</sup> |
| --- | --- |
| Minimal<sup>a</sup> | Tryptophan, cysteine, isoleucine, leucine, methionine, proline, valine, R-pantothenate, Pyridoxine, biotin, glucose |
| BDM<sup>b</sup> | Minimal plus thiamine, nicotinamide, Riboflavin, 4-aminobenzoate, folic acid, vitamin B12 |
| CADM<sup>c</sup> | BDM plus the 13 standard amino acids not listed in the minimal medium |
| Complex<sup>d</sup> | All the components of the minimal medium, the remaining 13 standard amino acids, riboflavin, folic acid, inosine, vitamin B12, chorismate, glucose, glycerol-3-phosphate, ribose, hypoxanthine, deoxycytidine, uracil, uridine, dexoyadenosine, adenosine, thymidine |

<sup>a</sup> (Hafiz & Oakley, 1976)
<sup>b</sup> (Karasawa *et al*, 1995)
<sup>c</sup> (Yamakawa *et al*, 1994; Haslam *et al*, 1986)
<sup>d</sup> (Oh *et al*, 2007)
<sup>e</sup> The following metabolites or proteins are added in SA analysis to permit the production of biomass: ATP, nicotinate, oxidized ferredoxin, oxidized thioredoxin, acyl and sulfur carrier proteins and are present in unchanging equilibrium concentrations in FBA.

### Essential metabolites

In order to validate the network, experiments involving the removal or addition of certain metabolites from the media were reproduced *in silico*. Each essential amino acid (cysteine, leucine, Isoleucine, proline, tryptophan and valine) was confirmed essential in the network as their removal prevented the production of biomass in any medium. None of the three essential vitamins are essential in the network (Table E1). The essentiality of two of these, biotin and pyridoxine, is due to their implication in the regulation of processes which could not be simulated in the metabolic network. Panthothenate (which is important in lipid metabolism) was in the past determined to be essential for a number of *C. difficile* strains tested (Karasawa *et al*, 1995). More recently, a new ketopantoate reductase (KPR) gene panG was discovered in a number of pathogenic bacteria and found to have a homolog in *C. difficile* strain 630 (Miller *et al*, 2013). Therefore, whereas panthothenate is commonly thought to be essential, this essentiality is strain specific and absent in *C. difficile* strain 630. A ΔpanG mutant in *Francisella tularensis* did not have any differences compared to wild type infections in a mouse model for pneumonic tularemia (Miller *et al*, 2013). This is likely due to the fact that panthothenate (vitamin B5) is widely available in food and most bacteria are able to import panthothenate through a sodium co-transport mechanism (Miller *et al*, 2013).

### Non-essential metabolites

Removal of non-essential metabolites did not have an important effect on growth. The non-essential amino acid methionine is known to enhance growth of the bacteria and is used in the minimal medium to increase growth rate. Interestingly, the removal of methionine from the minimal medium leads to a slight reduction in biomass production (less than 1%), a small but qualitatively correct effect. We simulated the removal of arginine and histidine (both non-essential amino acids) from the rich medium and in both cases this lead to the qualitatively correct result of a decrease in biomass production in agreement with the experimental evidence (Hafiz & Oakley, 1976). Most (8 out of 11) non-essential amino acids that when removed from a complete media do not affect growth rate experimentally also have no effect *in silico* when removed from the complex medium. Furthermore, the addition of certain non-essential amino acids (7 out of 14) or any nucleosides to the minimal medium leads to an augmentation of biomass production in agreement with experimental data (Table E1).

### Carbon sources

We tested the effect of removal of glucose as carbon source in the minimal and complex media as well as the use of alternative carbon sources. The removal of glucose produced the largest decrease in biomass production (28%) other than full arrest observed in the network. In complex medium however the effect was much smaller (5.86%) where amino-acid catabolism can be used as a source of carbon. The utilisation of different carbon sources in the absence of glucose in the network was simulated and compared to experimental data. Such data is not always specific to *C. difficile* strain 630 and small differences among strains do exist (Hafiz & Oakley, 1976) (Table E2). For 15 experimentally tested carbon sources (out of 20 carbon sources tested *in silico*) we obtain 100% agreement between the predicted utilization of alternative carbon sources by *C. difficile*, including the impossibility to use lactose as a carbon source. Furthermore, we predict that *C. difficile* strain 630 should not be able to use rhamnose or myo-inositol (for which there is some evidence of usage in other strains but no experimental information for strain 630) while malate, glycerol and chorismate would lead to increased growth rates.

### Comparison with existing metabolic network reconstructions

We compared *i*MLTC804cdf to the recently created automatically-generated non-curated reconstructed metabolic network of *C. difficile* (Büchel *et al*, 2013). The automated network contains 3211 reactions, 1548 unique metabolites and 1337 genes resulting in over 2762 genes/reactions associations. One fundamental requirement for a reconstructed metabolic network is its ability to produce biomass. As noted by its creators, the automatic network cannot produce biomass. This is likely due to the numerous flaws present in the automated network that are absent or present in a lesser number in the curated *i*MLTC804cdf reconstruction presented here. Among these: generic metabolites, incorrect reaction stoichiometry, repetitions, unclear reactions, dead-end metabolites and non-metabolic genes and reactions. For example, the automated network contains 485 generic metabolites representing 31.3% of the metabolome opposed to 23 in *i*MLTC804cdf (representing 3.8% of the metabolome). Curiously the automated network reconstruction contains 29 reactions involving oxygen, which should not be present in an anaerobic organism. Other important flaws in the automatically generated *C. difficile* network include the presence of 1562 export reactions and 648 dead-end metabolites. Overall, only 532 reactions (58.9% of the reactions in *i*MLTC804cdf) are common to both reconstructions out of 3211 present in the automatically generated network. A detailed comparison is presented in Table II (or in the form of a Venn diagram in Figure E1) and clearly shows the vast differences between the two models and that a large number of problems associated to the automatically generated network are absent in *i*MLTC804cdf.

**Table II:** Comparison between the automatic *C. difficile*, *i*MLTC804cdf and *C. acetobutylicum* networks.

| Characteristic | Automatic network | iMLTC804cdf | *C. acetobutylicum* |
| --- | --- | --- | --- |
| Reactions | 3211 | 911 | 552 |
| With genes | 1311 (40.8%) | 707 (77.4%) | 414 (75.0%) |
| With an EC number | 1435 (44.7%) | 747 (81.9%) | 497 (90.0%) |
| With KEGG ID | 1038 (32.3%) | 702 (77.19%) | 442 (80.1%) |
| Identified as unclear reaction | 4 (0.4%) | 0 | 1 (0.2%) |
| Associated with multiple reactions | 84 (8.1.6%) | 6 (0.7%) | 0 |
| Involving polymers | 26 (0.8%) | 0 | 0 |
| Involving generic metabolites | 548 (29.8%) | 31 (3.4%) | 35 (6.3%) |
| Non-metabolic reactions | 138 (4.3%) | 2 | 2 |
| Involving oxygen | 29 (0.9%) | 0 | -<sup>b</sup> |
| Transport reactions | 1576 (49.1%) | 145 (15.9%) | 80 (14.5%) |
| Export | 1562 (99.1%) | 16 (11.0%) | 6 (7.5%) |
| Passive | 14 (0.9%) | 15 (10.3%) | 17 (21.3%) |
| with genes | 0 | 95 (65.5%) | 61 (76.3%) |
| Unique reactions in common | 532 |  | 435 |
| Metabolites | 3133 | 705 | 515 |
| Unique | 1551 | 593 | 462 |
| With KEGG ID | 1092 (70.4%) | 563 (94.9%) | 436 (94.4%) |
| Generic | 485 (31.3%) | 23 (3.8%) | 28 (6.1%) |
| Duplicated | 15 (1.0%) | 0 | 0 |
| Dead-end | 648 (41.8%) | 8 (1.2%) | 52 (11.3%) |
| Unique metabolites in common | 533 |  | 372 |
| Genes | 1336 | 804 | 475 |
| Unique loci | 788 | 804 | 475 |
| With a homolog<sup>a</sup> | 788 (100%) | 658 (81.8%) | 417 (87.8%) |
| Non-metabolic loci | 66 (8.4%) | 3 (0.4%) | 7 (1.5%) |
| Loci linked to multiple entries | 382 (48.5%) | 0 | 0 |
| Unique loci in common | 534 |  | 357 |

<sup>a</sup> The values in the first and last columns represent the number of homologs with respect to iMLTC804cdf. The values in the middle column represent the number of homologs with respect to *C. acetobutylicum*.
<sup>b</sup> Reactions involving oxygen are allowed in *C. acetobutylicum* thus were not quantified.

We also compared *i*MLTC804cdf to the curated metabolic network reconstruction of the closely related bacterium *Clostridium acetobutylicum* (Senger & Papoutsakis, 2008). The analysis of reactions shared between the two reconstructions was possible due to the extensive use of KEGG identifiers in both networks. In the case of transport reactions, two transporters were considered similar if they transported the same molecule with one transporter from one network potentially matching multiple transporters in the other network. We did not differentiate between phosphoenolpyruvate (PEP):carbohydrate phosphotransferase system (PTS) (Postma *et al*, 1993), ion channels (Delcour, 2013) or ATP driven transporters (Hollenstein *et al*, 2007) as long as the transported molecules were the same in the two networks. The *C. acetobutylicum* network presents certain irregularities (misspelled metabolite names) that were corrected before the analysis. The network of *C. acetobutylicum* can only be analyzed using the proprietary LINDO API (Lindo Systems, Chicago, IL) which is paid software not optimized for use on biological networks (e.g. no specific functions for gene deletion studies). The absence of a commonly-accepted usable version of the network such as SBML (Hucka *et al*, 2003) or TSV (Gelius-Dietrich *et al*, 2013) both usable with R or Matlab, prevented a deeper analysis of the network. *i*MLTC804cdf contains 359 more reactions than the reconstructed network of *C. acetobutylicum* and 436 in common with it (representing 77% of its reactions). *i*MLTC804cdf contains 131 extra unique metabolites and 372 metabolites in common (representing 81% of its unique metabolites). The presence of 52 dead-ends metabolites (11.3% of the metabolome) suggests that the *C. acetobutylicum* network still has incomplete pathways and gaps that could affect biomass production. A detailed comparison is presented in Table II or in the form of a Venn diagram in Figure E2.

### Single gene deletions

We performed an *in silico* gene deletion study using both Synthetic Accessibility (SA) and Flux Balance Analysis (FBA) on *i*MLTC804cdf in order to identify potential essential genes that may lead to the discovery of novel therapeutic targets. This analysis removed reactions that were catalyzed by each gene alone (or in pairs, next section) or by a complex that involved that gene product and then measured the capacity of the network to produce biomass (either a flux in biomass production for FBA or S_net_ for SA, see material and methods). The complex medium (as describe in Table I) was the one used for the gene deletion studies since it reproduces the high concentration and diversity of nutriments found in the intestinal tract. Furthermore, the medium used is the same as that used in the simulation of the *Bacillus subtilis* metabolic network (Oh *et al*, 2007), which in turn is an approximation of the one used to perform the experimental validation of lethality of single gene deletions in that organism (Kobayashi *et al*, 2003).

A total of 73 out of 804 genes deletions were identified via FBA analysis as deleterious based on a 5% variation threshold (Shlomi *et al*, 2008). An additional 10 genes were found by SA to increase the number of reactions necessary to produce biomass and deemed essential based on this criterion. Overall, 50 of the 83 predicted essentials genes are essential according to both FBA and SA (see Table III). We observe an agreement rate of 95.8% between the two techniques in terms of the prediction of both lethal and non-lethal genes, which is similar to what was found when comparing both techniques on *E. coli* and *C. cerevisiae* (Wunderlich & Mirny, 2006).

**Table III.** List of 83 predicted *C. difficile* essential gene and essentiality of their *B. subtilis* homologs.

| Essentiality of *B. subtilis* homolog | Predicted *C. difficile* essential genes<sup>a</sup> |
| --- | --- |
| Essential (48) | accA<sup>3</sup>, accB<sup>3</sup>, accC<sup>3</sup>, accD<sup>3</sup>, **acpS**<sup>2</sup>, **adk**<sup>2</sup>, **asd**<sup>3</sup>, cdsA<sup>3</sup>, cmk<sup>2</sup>, **coaE**<sup>3</sup>, **dapF**<sup>3</sup>, **dapH**<sup>3</sup>, **ddl**<sup>3</sup>, **dxr**<sup>3</sup>, **dxs**<sup>3</sup>, **fabD**<sup>3</sup>, **fabF**<sup>3</sup>, glmU<sup>3</sup>, gmk<sup>3</sup>, guaB<sup>2</sup>, **ispD**<sup>3</sup>, **ispE**<sup>3</sup>, **ispF**<sup>3</sup>, **ispG**<sup>3</sup>, **ispH**<sup>3</sup>, metK<sup>2</sup>, mnaA<sup>3</sup>, **murA**<sup>3</sup>, **murB**<sup>3</sup>, **murC**<sup>3</sup>, **murD**<sup>3</sup>, **murE**<sup>3</sup>, **murF**<sup>3</sup>, **murG**<sup>3</sup>, nadD<sup>3</sup>, **nadE**<sup>3</sup>, **pgsA**<sup>3</sup>, plsX<sup>3</sup>, prs<sup>3</sup>, **CD0119**<sup>3</sup>, **CD0244**<sup>3*</sup>, CD1049<sup>3</sup>, CD2439<sup>3</sup>, CD3543<sup>2</sup>, CD3550<sup>3*</sup>, pgk<sup>1</sup> (15.48%), fchA<sup>1*</sup> (10.25%), folD<sup>1</sup> (10.25%) |
| Non-essential (26) | coaBC<sup>3</sup>, coaD<sup>3</sup>, **crtI**<sup>3</sup>, deoB<sup>1</sup>, deoC<sup>1</sup>, **fabH**<sup>3</sup>, **fabZ**<sup>3</sup>, guaA<sup>3</sup>, ispA<sup>3</sup>, metF<sup>1*</sup>, pdp<sup>3*</sup>, **pheA**<sup>1</sup>, **ribC**<sup>3</sup>, **tyrC**<sup>1*</sup>, **CD0557**<sup>2</sup>, CD1966<sup>3*</sup>, **CD2256**<sup>2*</sup>, CD2541<sup>2*</sup>, **CD2549**<sup>2*</sup>, CD3555<sup>3</sup>, CD3596<sup>1</sup>, pykF<sup>1</sup> (15.51%), **CD0994**<sup>1*</sup> (15.08%), **serA**<sup>1</sup> (15.08%), **ackA**<sup>1</sup> (14.74%), **asnA**<sup>1</sup> (5.76%) |
| No *B. subtilis* homolog (9) | **catI**<sup>1*</sup>, ntpA<sup>1*</sup> (10.31%), ntpB<sup>1</sup> (10.31%), ntpC<sup>1</sup> (10.31%), ntpD<sup>1*</sup> (10.31%), ntpE<sup>1*</sup> (10.31%), ntpF<sup>1*</sup> (10.31%), ntpK<sup>1*</sup> (10.31%), fhs<sup>1*</sup> (10.25%) |

<sup>a</sup> Number in parenthesis represent the loss of biomass according to FBA. Those without number prevent the production of biomass or were identified only by SA. Genes in bold (27 in total) do not have a human functional homolog. The 4 underlined genes have a human homolog shown to be essential when their associated reactions are removed from RECON2. Genes in green and red respectively were shown to be up- or down-regulated *in vivo* (Janoir *et al*, 2013). Genes marked with an asterisk (18 in total) are not present in the Database of Essential Genes (DEG) (Luo *et al*, 2014).
<sup>1</sup> Genes predicted as essential exclusively by FBA analysis (23 in total).
<sup>2</sup> Genes predicted as essentials by the SA analysis (10 in total).
<sup>3</sup> Genes predicted as essentials by both FBA and SA (50 in total)

The utilisation of two different techniques for *in silico* gene deletion provided a better coverage of the ensemble of genes that could potentially have an effect on the metabolism. The usage of SA allowed the identification of targets such as acpS, which is the only gene that can catalyze the transition of an apo acyl-carrier protein to a holo acyl-carrier-protein that participates in the elongation of lipids. In FBA this reaction is not required since a cycle is formed to reuse the holo acyl-carrier protein released at the end of the elongation and thus has no effect on flux in equilibrium. Using this cycle FBA can remain in a steady state and produce biomass but in the bacteria, production of the protein is required and acpS is likely to be essential as it is in many species (10 species according to DEG, the Database of Essential Genes (Luo *et al*, 2014)).

Essential genes were compared to experimental results for the gram-positive bacteria *Bacillus subtilis* (Kobayashi *et al*, 2003), which is the closest relative of *Clostridium difficile* with experimental essentiality data for all of its genes. Since the simulation involved the deletion of genes via deletion of metabolic reactions, functional homologs (genes responsible for reactions that share the same EC number and catalyze similar reactions) were used for the comparison. Among the 83 genes with a predicted effect on biomass production in *C. difficile*, 48 have homologs that are essential in *B. subtilis*, 9 did not have any functional homolog and 26 had a homolog that was not essential in *B. subtilis* (Table 3). We were able to compare 615 *C. difficile* genes (76.5% of the genes in *i*MLTC804cdf) for which we could detect a *B. subtilis* functional homolog with an overall prediction accuracy of 88.8% (Table E3). A similar rate of 89.0% was obtained based on the comparison of 520 genes using sequence homology (E value < 1e^−5^, sequence identity above 30% and alignment overlap over 80% of the *C. difficile* sequence, Table E3). While an accuracy of around 89% is extremely high, it is important to keep in mind that despite being closely related, differences are expected between the *B. subtilis* and *C. difficile*.

In order to compare our predicted essential genes with those predicted to be essential in *C. acetobutylicum* (Senger & Papoutsakis, 2008), we selected the subset of 357 genes in *C. difficile* with a sequence homolog in the *C. acetobutylicum* network. Of these, 127 genes reproduced the effect of a single reaction removal in our network and could be compared with the reaction removal study above performed by Senger & Papoutsakis. This comparison was made using slightly different media for each bacterium both described either as complex or minimal to ensure that the media had the specific elements required by each species. We obtain an agreement of 76.38% in complex media and 68.50% in minimal media as shown in Table E4.

The inhibition of the product of essential genes that are upregulated during CDI may require a smaller drug dose to generate an effective response, thus decreasing side effects. We utilized transcriptomics data associated to the differential expression of genes during infection (Janoir *et al*, 2013) to annotate predicted essential genes in view of their use potential use as therapeutic targets. Eight predicted essential genes are downregulated *in vivo* while 6 are upregulated during infection. Some of the genes that are upregulated during infection and predicted to be essential such as fabD, serA or CD2549, are of additional interest as they could not only affect growth, but also colonisation and pathogenesis processes (Janoir *et al*, 2013) (Table III).

We compared the list of 83 genes predicted to be essential in *C. difficile* using *i*MLTC804cdf to the genes in the Database of Essential Genes (DEG) (Luo *et al*, 2014). Interestingly, and serving as further validation of *i*MLTC804cdf, a total of 65 of these genes are present in DEG, i.e., these genes have homologs known to be essential in other species. The remaining 18 predicted essential genes that are not present in DEG (Table III) include ntpA,D,E &F, K (all subunits V-type ATP synthase) as well as tyrC, metF (involved in amino acid synthesis) and xpt (guanine synthesis) among others. Finally, we analysed the list 7 *C. difficile* potential drug targets present in the *C. difficile* Drug Target Database (clostridium-DTDB) (Jadhav *et al*, 2013). These 7 potential targets are originally derived from DEG but fulfill a number of additional conditions. Among the targets in clostridium-DTDB, we find a number of targets that have non-metabolic roles (thrB and cheR), that are not essential in *i*MLTC804cdf due to the existence of alternative pathways or isoenzymes (e.g., nanE and bioB) or for which the metabolic product of the reaction is supplied in rich media (e.g., pyrF and panC). Only deoB, present in clostridium-DTDB as a potential target is also essential in *i*MLTC804cdf (Table E5).

Lastly, despite the lack of extensive information on experimentally verified essential genes, the little evidence that exists, supports our predicted essential roles for a number of genes: metK and trpS (Walker, 2012), guaA (Mulhbacher *et al*, 2010) and ntpA-B-C-D-E-F-K (Wu *et al*, 2013). In one case, CD0274 (dhaT in *i*MLTC804cdf, also called gldA), while the gene was shown to be essential, this effect is not due to a metabolic role but rather due to the detoxification of metabolic byproducts (Liyanage *et al*, 2000). Other known essential genes such as secA1-A2 (Fagan & Fairweather, 2011), metG and gyrA-B (Oh & Edlund, 2003) are not present in the network and are involved in non-metabolic processes. Lastly, the gene prdF has been mutated and was shown to be non-essential (Wu & Hurdle, 2014), in agreement with its predicted non-essential role in *i*MLTC804cdf.

### Detection of potential human cross-reactivity targets

#### Sequence and functional similarities

One of the main goals in detecting essential genes is to assess their potential as therapeutic targets. One factor weighting in favour of a potential therapeutic target is the lack of a human homolog, as this decreases the chances of side effects of a potential drug off-targeting the gene product of the human homolog. We again utilize here two definitions of homology, the standard sequence homology that relates two genes through evolution and functional homology, that relates two genes via common function of their gene products, specifically the same E.C. number. Functional homology is stricter than sequence homology as it is not based on any level of similarity between the two proteins, only based on the fact that the two enzymes catalyze the same reaction. Twenty-seven genes identified as potential targets do not have any human functional homolog (58 based on sequence homology). In other words, 56 genes have functional human homologs and 25 have sequence human homologs out of the 83 predicted essential *C. difficile* genes (Table E5).

#### Metabolic essentiality

In the absence of experimental evidence of the potential for the cross-inhibition of the human homolog by a potential drug, we sought to use FBA to determine if inhibition of the human homolog would have any effect on the human cell. To do so, we performed a gene deletion FBA analysis on the latest draft of the human reconstructed metabolic network RECON2 (Thiele *et al*, 2013). In the case of functional homologs, we opted for a conservative approach in which all reactions associated to the human homolog of the *C. difficile* target were removed for the FBA analysis from the human network. Thus simulating a situation in which a potential drug would inhibit all potential cross-reactivity targets. In total, only 4 predicted essential *C. difficile* genes out of 49 tested have predicted essential human homologs. For the remaining 34 genes, 8 have human homologs that were not present in RECON2 and 26 have no human functional homologs. This analysis suggests that the presence of a human homolog may not be sufficient to discard a potential target due to the possibility that an eventual drug interacts with the human protein. This is due to the fact that the inhibition of the human homolog may not have any serious effect in human cells given the differences between the two metabolic networks.

#### Binding-site structural similarities

We proceeded to utilize the detection of binding site similarities to detect potential cross-reactivity targets for the 26 *C. difficile* proteins where sequence or functional homology did not detect any human homolog. To perform this analysis, we created homology models of the targets using I-Tasser (Roy *et al*, 2010). The best model generated by I-Tasser for each protein was submitted to analysis using IsoCleft (Najmanovich *et al*, 2008; Kurbatova *et al*, 2013), a method to detect local 3D atomic similarities between binding-sites. In each case the largest cleft (representing the binding site in 83% of cases (Laskowski *et al*, 1996)) identified by IsoCleft was searched against a non-redundant dataset of 7339 binding-sites of unique combinations of protein families bound to distinct ligands (Kurbatova *et al*, 2013). In each case we selected the most similar match found to a protein in the IsoCleft Finder non-redundant dataset that contains human homologs. It is important to note however that the representative of such family in the non-redundant dataset may itself not be necessarily a human protein. For 23 out of 26 models we found on average 37 atoms in common (average p-value 0.038 and z-score 3.05) with members of Pfam families (Punta *et al*, 2011) that include human proteins. In three cases, ispF, CD2549 and dapH, no significant level of binding-site similarity was found to a Pfam family that contains human homologs (Table E6).

It is hard to judge if matches found are significant or not considering that no threshold for binding site similarity can be uniquely defined (Najmanovich *et al*, 2008) above which cross-reactivity is certain. However, in 18 cases out of 26, the top-scoring detected binding-site similarities for each case represent binding-sites in proteins that bind ligands that are similar to at least one of the substrates of the reaction catalyzed by the modelled *C. difficile* protein. In seven of those cases, the top matching Pfam family that contains human homologs binds a similar ligand (Table E6). Taking as an example the case of the enzyme encoded by the asd gene, we detect 39 atoms (Z-score 3.92, p-value 0.012) in common to a glyceraldehyde-phosphate dehydrogenase from spinach (PDB ID 2PKQ) bound to NADPH, a member of Pfam family PF00044, that has human homologs (Figure 1). Five out of the top 7 most similar binding-sites also bind NADPH or NADP, all from different Pfam families. The superimposition of these diverse binding-sites based on their similarities to the asd gene product binding-site leads to an extremely good superposition of their respective bound-ligands (Inset Figure 1). This suggests that the detected similarities are biologically significant. The quality of the resulting superimpositions together with the detection of similarities across families that bind similar ligands to those that bind the *C. difficile* targets reinforces the confidence in the biological significance of our predictions. The quality of the alignment of the NADP molecules across different families via the detected similarities suggests that these capture the molecular determinants responsible for binding. As such determinants are conserved across different families, there is a possibility that these are also conserved within families and thus present in the human homolog. In most cases, the similarities that were detected actually represent commonly used cofactors or other ubiquitously used ligands, such as NADP above or ATP. These results don’t necessarily mean that a drug targeting the *C. difficile* protein will bind the human homolog belonging to the detected Pfam families, but these should be used as potential cross-reactivity targets in the rational design of inhibitors against the *C. difficile* protein in question. Furthermore, given that the detected similarities focus on common cofactors and ubiquitous molecules such as ATP, the results also suggest that targeting the sub-pockets of less ubiquitously used substrates may reduce the chance of cross-reactivity.

**Figure 1:**
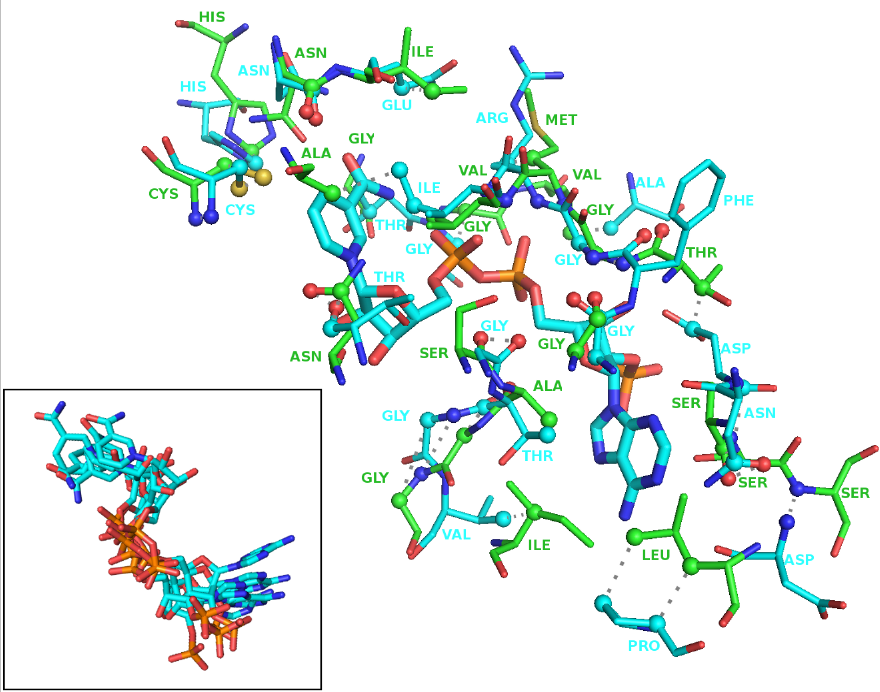
Example of biding site similarities between the modelled asd gene product and the photosynthetic a2b2-glyceraldehyde-phosphate dehydrogenase bound to NADP. The two binding-sites share 39 atoms of equivalent atom types in corresponding positions in space (Z-score 3.92, p-value 0.012). This protein from spinach (PDB ID 2PKQ) belongs to Pfam family PF00044 that contain human homologs. The inset shows the superimposition of the bound NADP molecules found among 5 of the top 7 most similar binding-sites belonging to different protein families. Carbon atoms in residues belonging the query asd protein cavity are marked in green while those in the target 2PKQ structure as well as those in the bound NADP are shown in cyan. Detected equivalent atoms are shown as spheres and grey dashed lines highlight pairwise equivalences.

### Double gene deletions

We performed double gene deletions to identify potential polypharmacological targets and to target reactions that are catalyzed by isoenzymes. Based on FBA analysis, 151 gene pairs involving 63 unique genes that had small or no effect in single gene deletion were deleterious when removed in pairs. An additional 3 essential gene pairs involving 5 new unique genes were found using SA. Eight gene pairs were considered essential in both SA and FBA analysis (Table E7). Some double mutants show a synergistic effect, defined as an effect greater that an additional 1% reduction in biomass production in the double mutant than the sum of effects of each single mutant.

The 68 synergistic double mutants were analysed in more detail (Table E8). Eleven of these synergistic gene combinations resulted in total abolition of biomass production in FBA or prevented the biosynthesis of at least one element of the biomass in SA, 4 had an effect between 10% and 20% and the remaining 53 had a small effect on biomass production (between 5 and 10%). Fourteen of the essential pairs of genes are isoenzymes that catalyze the same reactions. Three gene pairs represent enzymes involved in pathways with the same functional category while the remaining 51 gene pairs affect different pathways (Figure 2). All essential pairs identified by both SA and FBA are isoenzymes whose removal results in a total arrest of biomass production.

**Figure 2:**
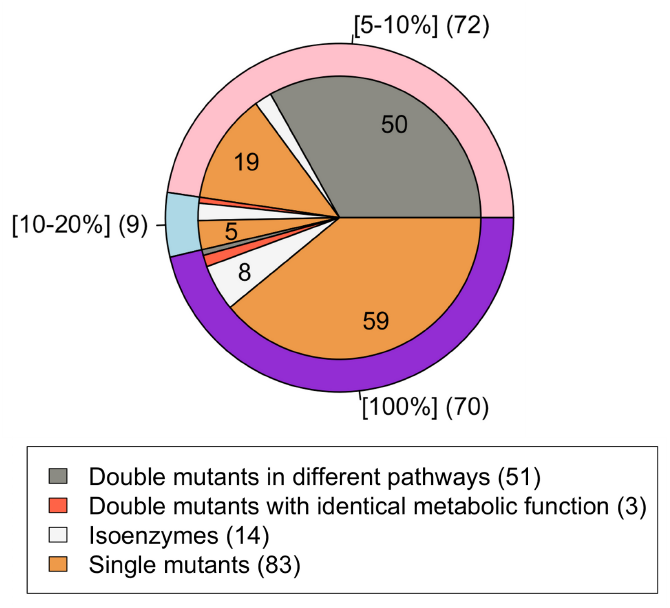
Effect of the deletion of essential genes and deletion of essential pairs of genes in term of biomass lost. The numbers in brackets represent the extent of decrease in biomass production. The number of cases in the unlabelled sections of the pie chart is in clockwise order 2, 1, 3, 1 and 3. In the case of SA predicted genes, were the effect on biomass production cannot be quantified beyond full arrest (100% decrease), the percentage decrease in biomass production is taken to be inverse the percentage increase in the number of reactions necessary to produce biomass.

Related-genes (isoenzymes or genes involved in the same metabolic function) usually result in a higher biomass loss than relatively distant pairs (Figure 2). For the 14 isoenzymes, the deletion of the two genes in a pair is required to remove a reaction that is catalysed by both. For the remaining essential gene pairs, the reactions associated with both genes are used in parallel in the wild type, a case of metabolic plasticity (Güell *et al*, 2014), or the reactions from only one of the genes is used while the reaction from the other member of the pair can act as a backup, a case of metabolic redundancy (Güell *et al*, 2014). Depending on the category in which an essential gene pair falls, different strategies may be required in order to target the pair (Güell *et al*, 2014). From the 54 essential genes pairs, 35 represent cases of plasticity of the network and 19 cases of redundancy in the network (Table E8).

### Distribution of predicted essential genes across pathways

In order to see which parts of metabolism are more enriched with essential genes, we classified reactions into 8 functional pathway classifications (Table E9). This analysis confirmed that linear pathways, like lipid synthesis, tend to have more essential genes (25 out of 74 genes in lipid synthesis pathways are essential) due to the lack of alternative ways leading to the production of highly important metabolites in biomass composition (Figure 3).

**Figure 3:**
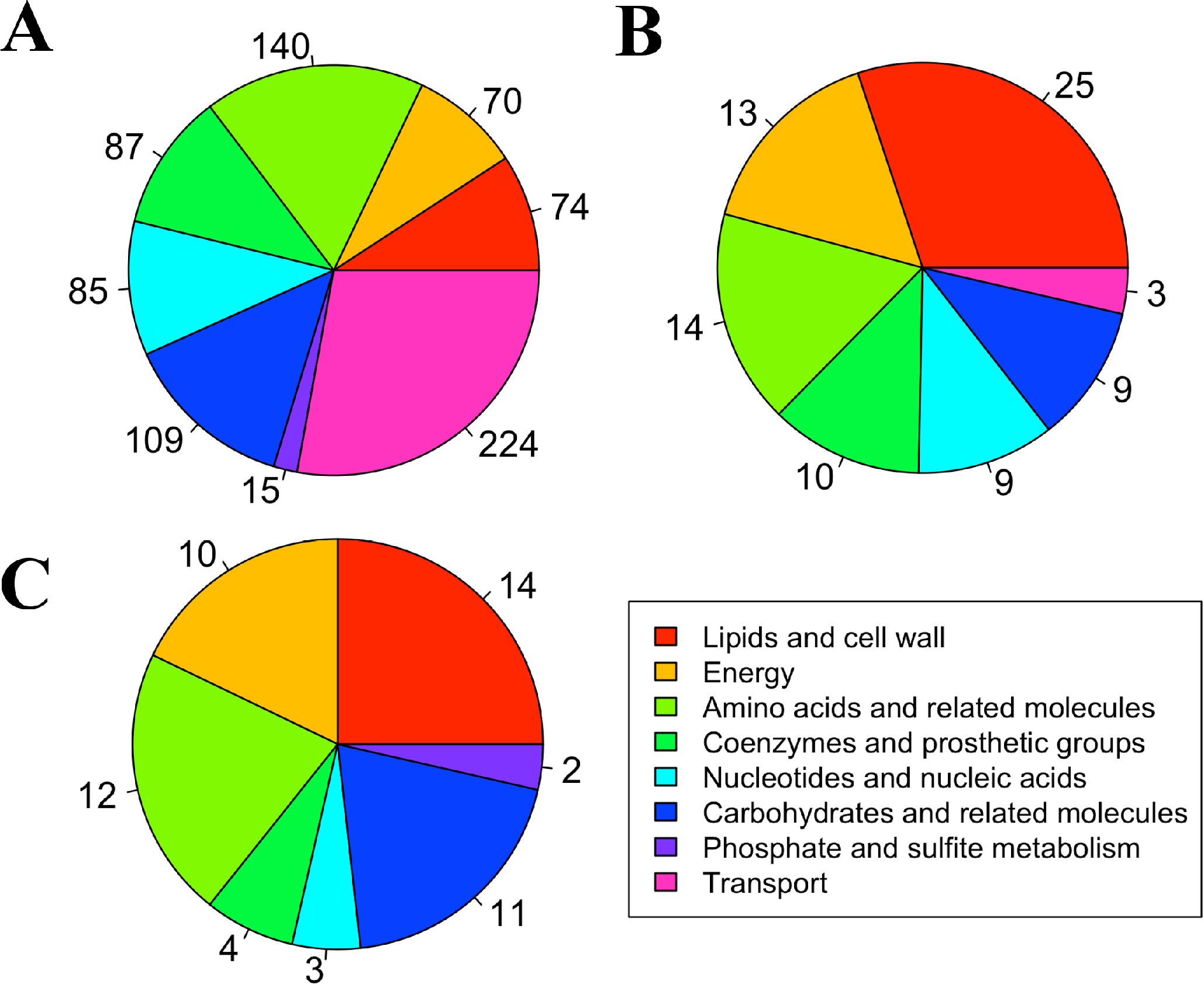
Distribution of genes (A), essential genes (B) and essential genes gene pairs (C) in the different metabolic pathways. If one gene was present in more than one reaction that did not share the same metabolic function the most relevant function was manually chosen based on gene description. We observe an enrichment of essential genes involved in lipid and cell wall production and a depletion of predicted essential genes involved in transport.

### Potential compounds binding predicted essential targets

Molecules potentially binding the proteins encoded by the 133 genes identified as potential targets on their own (83 genes) or as part of pairs (50 unique genes, forming 68 different pairs with deleterious effect when both genes are removed) were identified based on sequence homology (E-value<1e^−5^, sequence identity above 30%, and overlap over 80%) between predicted essential proteins and entries in the DrugBank database Version 4 beta (Law *et al*, 2014) (Table IV). A total of 72 molecules bind 38 protein entries from DrugBank with homologs among 19 predicted essential *C. difficile* targets. While the list includes cofactors, binders, inhibitors and activators, all such molecules bind the homologs of the predicted essential *C. difficile* targets. Most of these molecules are still experimental. Interestingly, 9 molecules are approved drugs based on DrugBank annotation. Among these we have Celurelin predicted to bind to the product of two distinct predicted essential genes in the same pathway: fabF and fabH (with the potential for polypharmacology). Mycophenolic acid and Ribavirin are predicted to bind the predicted essential product of guaB. Tretinoin and 2PP both inhibitors of the product of gapN which is essential only as part of double mutant with different subunits of the F-type ATP synthases (atpA,B,D,F,G,H,I,Z). The double inhibition could be achieved with the use of any drugs that would affect ATP production such as nisin, reutericyclin-867 or valinomycin (not identified via sequence identity) (Wu *et al*, 2013). Lastly, pyridoxal phosphate is a potential binder of two proteins that are part of pair whose double mutation is predicted to be lethal (glyA and CD2834). The identification of any potential binding small-molecules (based on target homology) is useful since there is a chance that these molecules may also bind a predicted essential *C. difficile* homolog protein and this information could be used as a basis for the development of specific inhibitors. This analysis also helps elucidate the role of pyridoxine, an essential vitamin that has no direct effect on biomass, since pyridoxal phosphate is a cofactor that binds two genes (glyA and CD2834) part of the same essential pair. This list remains to be experimentally validated but is meant as a starting point in targeting one of the genes predicted as being essential in the network.

**Table IV.** List of 19 potential targets associated with 72 potential binders based on sequence identity.

| Target (pair)<sup>a</sup> | Sequence identity<sup>b</sup> |  | DrugBank Potential inhibitors<sup>c</sup> |
| --- | --- | --- | --- |
|  | *E.coli* | Human |  |
| accC | P24182 [53%] | Q96RQ3 [48%] | LZJ, LZK, LZL, OA1, OA2, OA3, OA4, OA5, L21, L22, L23 (P24182 [53%]); BTI (Q99UY8 [48%]); D1L, H1L (O00763 [32%]) |
| adk | P69441 [52%] | P54819-2 [45%] | <u>GEO</u> (P30085 [32%]) |
| fabF | P0A6Q6 [49%] | Q9NWU1 [45%] | <u>CER</u>, P9A, PMN (P0AAI5 [50%]); TL5, TL6, P4T (P0A953 [38%]); TLM (P63454 [37%]) |
| fabH | P0ACC7 [42%] | - | <u>CER</u> (P0A6R0 [45%]); B82 (820T1 [43%]); UDT, VZZ, D1T, DFD, MDX (P0A574 [37%]) |
| fabZ | P0ADG7 [36%] | P20839 [35%] | QUE, 2BC, 2BE, 2RB, 3BE, 4BB, 4BE, AGI, EMO, SAK (Q5G940 [41%]); BDE (O25928 [41%]), |
| glmU | P62617 [55%] | - | P21 (P43889 [42%]) |
| guaB | P14900 [32%] | P20839 [35%] | <u>MOA</u>, <u>RBV</u> (P20839 [35%]) |
| ispF | P0AD61 [51%] | P14618-2 [44%] | FPP (Q8EBR3 [62%]); GPP (P62617 [55%]); IPP (A0R559, [34%]) |
| murD | P0A858 [44%] | P60174-1 [44%] | LK1, LK2, LK3, LK4, LKM (P14900 [32%]) |
| pykF | - | - | D8G, DYY, DZG (P14618 [44%]) |
| tyrC | P0A6P9 [64%] | P09104 [52%] | HPP, ENO (O67636 [32%]) |
| fabG (CD2577) | P33898 [61%] | P04406 [44%] | EMO (P16544 [37%]); TAQ, CB3, DVP, FE1 (P0CG22 [34%]); TDB (Q16698 [31%])BRZ (O14556 [44%]); AES (P04406 [44%]) |
| gapA (gapB) | P0A9B2 [52%] | O14556 [47%] | BRZ (O14556 [44%]); AES (P04406 [44%])AES (P04406 [49%]); BRZ (O14556 [47%]) |
| gapB (gapA) | P25526 [35%] | P00352 [35%] | AES (P04406 [49%]); BRZ (O14556 [47%])<u>REA</u> (P00352 [35%]); DXC (P25554 [31%]); <u>2PP</u> (P51649 [31%]) |
| gapN (atpA-Z)<sup>d</sup> | P10902 [40%] | D6RFM5 [32%] | <u>REA</u> (P00352 [35%]); DXC (P25554 [31%]); <u>2PP</u> (P51649 [31%])UQ1, BHG (P31040 [32%]); HQO, BRS (P00363 [32%]); UQ2, DNT (P0AC41 [31%]) |
| uppS (CD2762) | P0A825 [56%] | P34897-3 [45%] | B08, B28, B29, B76, FPP, GPP (P60472 [44%]); IPE (P60379 [43%])<u>PLP</u> (P34897 [45%]) |
| glyA (CD2834) | P0A825 [48%] | P34897-3 [41%] | <u>PLP</u> (P34897 [45%])<u>PLP</u> (P34897 [41%]) |
| CD2834 (glyA) | - | B7ZLW7 [31%] | <u>PLP</u> (P34897 [41%])<u>CNC</u>, <u>I2A</u> (Q99707 [31%]) |
| CD0727 (phnA; aldH)<sup>d</sup> | P24182 [53%] | Q96RQ3 [48%] | <u>CNC</u>, <u>I2A</u> (Q99707 [31%]) |

<sup>a</sup> Genes that are essential as part of a double mutant pair have the name of the partner gene identified in parenthesis.
<sup>b</sup> Uniprot ID of the closest sequence homolog.
<sup>c</sup> 3-letter PDB code of ligand that is a known inhibitor or binder of a sequence homolog with the Uniprot ID and the level of sequence identity given in parenthesis.
<sup>d</sup> The target gene is part of more than one essential pairs, one with each of the proteins in parenthesis.

## DISCUSSION

In this work we present *i*MLTC804cdf, a highly curated metabolic network of the nosocomial pathogen *Clostridium difficile* strain 630. This metabolic network is functional in the sense of being amenable to simulations using Flux Balance Analysis to measure biomass production in diverse types of media. *i*MLTC804cdf is available in SBML, TSV and Excel formats. The network is based on the aggregation of metabolic data present in databases, augmented with information from various sources of experimental data from the literature and manually curated in order to ensure a high quality of the resulting network.

Metabolic networks bring together information from various databases. Due to inaccuracies, contradictions and missing information in each database (Stobbe *et al*, 2013; Latendresse *et al*, 2012; Green & Karp, 2005), the non-curated network resulting from the simple aggregation of all these sources of information is often incomplete and includes a large number of errors, missing data and repetitions, which make the generated networks unusable (Büchel *et al*, 2013). Different databases will often have multiple identifiers for highly similar entries, which cause duplications. Surprisingly, this is true not only for reactions and metabolites where curation is more difficult but also for genes. Manual curation is essential to correct inaccuracies present in metabolic databases and particularly to fill gaps (missing reactions) necessary to remove dead-end metabolites to create a functional network. Manual curation is a time consuming, expensive and non-scalable process. Unfortunately, it is also indispensable at this time as can be attested by the 21 predicted essential genes (alone or as part of a double mutant) that do not even appear in the automatically generated *C. difficile* network. It is clear that the automated generation of functional metabolic networks is a worthwhile goal. However, at present automatically generated networks are not functional. At this time, such networks can only be used at best as a starting point prior to intensive manual curation. Their use as starting points however subtracts very little to the manual work required to produce a functional network.

In some sense, metabolic networks serve as a tool to aggregate all existing knowledge about the metabolism of an organism. Clearly, the more well studied an organism is, the more information exists to build such a network. *C. difficile* is an organism that is not well studied due to a number of factors. First, being a pathogen severely restricts the number of active researchers studying it given the experimental requirements in terms of biosafety. Second, *C. difficile* presents particular challenges that make genetic manipulations notoriously difficult. Until recently, genetic studies in *C. difficile* were restrained by the lack of efficient tools to inactivate specific genes. The recent development of a universal gene knockout system in clostridia has opened new possibilities and it is now somewhat easier to disrupt genes in a very specific and directed manner (Heap *et al*, 2007; 2010). This system called ClosTron is based on retargeting of the *Lactococcus lactis* Ll.LTRB group II intron so that upon transfer into *C. difficile* by conjugation, the intron integrates at a specific user-defined chromosomal site. With the advent of ClosTron our knowledge of the biology of *C. difficile* and the identification of essential genes is bound to increase but at the moment, there is a lack of *Clostridium difficile* specific literature and that limits the completeness and validation of the network. Some information such as biomass constituents and protein-protein interactions had to be extrapolated from other closely related species. The metabolic pathways involved in linking dead-end metabolites to the rest of the network also sometimes had to be extrapolated for lack of experimental data. No in-depth studies of the directionality of reactions, as it was done in *E. coli* (De Martino *et al*, 2012), or association between biomass production and predicted growth rate, as it was done in *B. subtilis* (Oh *et al*, 2007), exist due to the absence of metabolomics data and precise growth rate experiments for *C. difficile*. While information can be extrapolated from closely related organisms, fundamental differences still exist and may be a source of potential errors. Notwithstanding the existing limitations in creating and validating a *C. difficile* specific network, *i*MLTC804cdf is as complete or more than existing curated networks and accounts for all existing relevant experimental information. We hope that in addition to being a tool to aggregate existing knowledge, *i*MLTC804cdf will prove to be valuable as a nucleation point in developing our understanding of this important human pathogen, driving the generation of new experimental hypotheses.

The analysis of a metabolic network in isolation in the absence of other relevant processes present in the organism poses its own problems. For example, the effect of metabolites (such as pyridoxine and biotin) (Karasawa *et al*, 1995) involved in non-metabolic processes could not be simulated. Likewise, methionine added to minimal media greatly increases growth *in vitro* (Hafiz & Oakley, 1976) but the addition of methionine to the minimal media in *i*MLTC804cdf increases biomass by less than 1%. Methionine is mostly used in the bacteria as S-adenosyl methionine involved in the biosynthesis of cofactors and vitamins which are not directly involved in biomass synthesis and have an effect that cannot be simulated in the metabolic network (Rodionov *et al*, 2004). While the removal of methionine produced a qualitatively correct outcome, the loss of biomass in the network when removing methionine from the minimal medium is only due to the additional reactions required to produce enough methionine as required for biomass production.

The case of pantothenate, the only essential vitamin with clear metabolic effect, is unique as its essentiality is strain-dependant. A biosynthesis pathway for panthothenate was recently identified in *C. difficile* strain 630 (Miller *et al*, 2013) and is present in some other strains based on MetaCyc. As a result, this vitamin is a non-essential component of the growth medium based on our *in silico* analysis. This vitamin is however essential in a number of strains previously tested (Karasawa *et al*, 1995), which did not include either strain 630 or the others containing this pathway in MetaCyc.

The comparison with *C. acetobutylicum* network (Senger & Papoutsakis, 2008) indicates that both bacteria share the same metabolic core. Existing differences in reactions and associated genes can explain the differences obtained while comparing the effects of deleted genes and reactions. The different media utilized for both bacteria can also explain part of the variation since the complex media for *C. acetobutylicum* is more complete then the one used for *C. difficile*. Deletions performed in minimal media increase the number of predicted essential genes but also make more evident specific differences between the organisms since they don’t share the same essential components (amino-acids, vitamins, *etc*.) (Monot *et al*, 1982; Hafiz & Oakley, 1976).

The gene deletion analysis identified interesting potential therapeutic targets. Targets such as the aspartate-semialdehyde dehydrogenase (E.C. 1.2.1.11) asd (UniProt ID Q17ZW9) or the diaminopimelate epimerase (E.C. 5.1.1.7) dapF (UniProt ID Q182T1, also known to be essential in *B. subtilis*) that don’t have human functional homologs, decrease the chance of side effects due to cross-reactivity. For these proteins binding-site similarities do detect potential cross-reactivity targets that should be considered as part of a rational drug design program. Another predicted essential gene, the aspartate-ammonia ligase (E.C. 6.3.1.1) asnA (UniProt ID Q183C9) is up-regulated *in vivo* and could be important for the pathogenesis of the bacteria (Janoir *et al*, 2013). Most targets like CD2549, dxr and fabD have more than one of these characteristics and would be interesting for more than one reason.

As a result of the conservation of local binding site environments (Kurbatova *et al*, 2013; Najmanovich *et al*, 2008; 2005), drugs often targets proteins in a way that might not be sequence-dependent. To account for that effect, we used functional homologs to identify potential human cross-reactivity targets for predicted essential *C. difficile* proteins. This made for a more stringent analysis since the number of potential human functional homologs is almost twice as large as those based on sequence similarity alone. The absence of a human homolog is often used as a criteria for identification of potential drug target (Jadhav *et al*, 2013). If no homolog is present, there is a smaller probability that a drug targeting this specific protein have an effect in humans.

The presence of a human homolog however is not sufficient to evaluate whether or not targeting a particular protein might have important side effects since the homolog might not be essential in humans. Targeting a *C. difficile* gene with a non-essential human homolog could still result in little or no side effects even if the human homologs are inhibited. The use of predicted essentiality of human functional homologs in RECON2 (Thiele *et al*, 2013) in conjunction with their essentiality in *C. difficile* represents a more consistent analysis of targets across hosts and pathogens. To our knowledge the use of functional homologs across species to determine the potential of a target to have cross-reactivity targets leading to side effects is novel.

For those cases where sequence or functional homology did not detect potential human cross-reactivity targets, we utilized the detection of local binding site similarities. This analysis identified protein families with human representatives that harbour large binding-site similarities to the *C. difficile* targets in the absence of sequence or functional similarities. The detected similarities are primarily localized to binding-sites of cofactors and ubiquitously used ligands such as NADP or ATP. Three genes, ispF, CD2549 and dapH, in addition to not having any human sequence or functional homologs, also show very little binding-site similarities to any protein family present in humans.

It is important to keep in mind that it is not possible to determine a minimum similarity threshold other than 100% above which one can be certain that the detected human proteins will act as cross-reactivity targets as small differences can bring about drastic effects (Najmanovich *et al*, 2008).

A “perfect” predicted essential target would be one with no or non-essential sequence, functional or structural homologs in human and *E. coli* (as a proxy for gram-negative and gut flora in general), with essential homologs in *B. subtilis* and up-regulated *in vivo*. Although no *C. difficile* target could be found fulfilling all properties at once, the 133 potentially essential targets identified (as single or double mutants) fulfil several of these properties and could, once validated experimentally serve as a target for the development of new antibiotics. New inhibitors should be designed to maximize the interactions with the *C. difficile* protein and minimize interactions with the detected potential human homologs.

The comparison with experimental results for *B. subtillis* (Kobayashi *et al*, 2003) was used as validation due to the absence of experimental results for the gene essentiality in *C. difficile*. While essential differences exist between the two organisms, a large degree of conservation is also present. Therefore one should expect that a large number of genes conserve their essentiality across these two species. The high level of accuracy (according to functional or sequence homology) between the experimental results and our predictions serves as a validation of *i*MLTC804cdf as a mature draft metabolic network and increases our confidence in the list of predicted essential genes. One important difference between the metabolism of *C. difficile* and *B. subtilis* is that the later can use oxygen to produce energy while the former cannot. Therefore, predicted essential *C. difficile* genes involved in fermentation such as pykF or ackA are not essential in *B. subtilis*. Other genes such as fabH, CD1966 and ribC are only present in one copy in *C. difficile* while more than one gene catalyses the same reactions in *B. subtilis* (Karp *et al*, 2005), explaining why such genes are non-essential in the latter.

Some genes whose inactivation is deleterious *in vitro* are not identified *in silico* due to their implication in non-metabolic processes. Both *i*MLTC804cdf and the network of *Bacillus subtilis* (Oh *et al*, 2007) fail to identify the essentiality of CD1536 (yumC in *B. subtilis*) due to its implication in regulatory processes. In addition, the toxicity of a molecule cannot be simulated *in silico* either. Therefore, the essentiality of genes involved in detoxification or whose deletion leads to the accumulation of toxic molecules cannot be simulated. For example, the removal of CD3543 would lead to an accumulation of nicotinate that could be toxic. This effect cannot be simulated in the network, therefore CD3543 is not considered essential in the network but is essential *in vivo* in *B. subtilis* (Kobayashi *et al*, 2003).

The combined use of FBA and SA allowed us to detect more essential genes than using either technique alone. Their joint use increases our confidence on the predictions for those genes where the two techniques agree. At the same time, the two techniques complement each other. For example, the gene acpS catalyses the only reaction that leads to the production of a holo-acyl-carrier protein from the apo version of the protein. This reaction is essential since the holo form of the protein is required to perform the elongation of lipids. The presence of a cycle that allows for the reutilisation of the released holo-acyl-carrier protein at the end of lipid elongation prevents FBA to identify the deletion of acpS as deleterious. The analysis by SA uses the apo version of the acyl-carrier protein and predicts acpS as essential since without it, the holo form cannot be produced. New targets were also found when using SA for double mutants. Targets identified by SA are mostly isoenzymes that lead to deletion of new reactions in the shortest possible pathway leading to the production of essential biomass metabolites.

The list of active molecules that potentially bind predicted essential targets includes many molecules that could help in the validation of the targets in *C. difficile* and the development of novel drugs (Shen *et al*, 2010). Experimental validation is required to determine if the identified small-molecules do bind the *C. difficile* homologs. Some of these small-molecules, such as the approved anti-viral Ribavirin, could speed the approval of *C. difficile* specific inhibitors through drug repositioning. In the case of Ribavirin, the molecule is a rapidly absorbed guanosine analog used in the treatment of Influenza (Sidwell *et al*, 2005) and hepatitis C (El-Shamy & Hotta, 2014). Cerulenin has anti-fungal and anti-bacterial activity targeting FabF in *B. subtilis* (Trajtenberg *et al*, 2014), thus very likely targeting the same protein in *C. difficile* as we predicted. In all cases, the multiple small molecules predicted to bind the predicted essential *C. difficile* proteins could be used to bias library selection for the rational development of new inhibitors.

## MATERIALS AND METHODS

### Creation and curation of the draft metabolic network

We created a collection of metabolic and transport reactions associated with *C. difficile* strain 630 from the KEGG (Kanehisa *et al*, 2014), MetaCyc (Caspi *et al*, 2014) and TransportDB (Ren *et al*, 2007) databases. Reactions involving polymers (glycogen, peptides and others) were ignored to avoid the spontaneous creation of matter due to the presence of molecules of undefined length. Also, some of these polymers, like glycogen, are only used for energy storage (Wilson *et al*, 2010) and wouldn’t have any impact in simulations that optimize biomass production. Reactions involved in spore formation, the conjugation process, RNA, peptide or DNA modification, cell repair, and other non-metabolic enzymatic reactions were not added to the network since these reactions aren’t directly contributing to the production of biomass constituents. Finally, the reconstructed network concentrates solely on metabolism without including signalling, gene regulation and post-translational modification of proteins even if these can clearly affect metabolism.

The initial draft network is little more than a collection of reactions. In that state it cannot be used for any sensible application such as the simulation of biomass production. This is due to the numerous inaccuracies present in the databases. These inaccuracies consist of, but are not limited to, the presence of generic and dead-end metabolites, missing or erroneous pathways, missing genes and unbalanced reactions (Table E10). Two cases exemplify some of these inaccuracies. First, noting that aerobic pathways should not be present at all in the *C. difficile*, the inclusion of incomplete versions of both aerobic and anaerobic pathways for the biosynthesis of vitamin B12 is problematic (Figure E3). Second, the omission of most reactions in the Stickland and amino acid fermentation pathways (Figure E4), important sources of energy for the bacteria, which had to be completed based on literature. Both of these corrections a numerous others in of the same nature were necessary to create a functional network (Figure 4).

**Figure 4:**
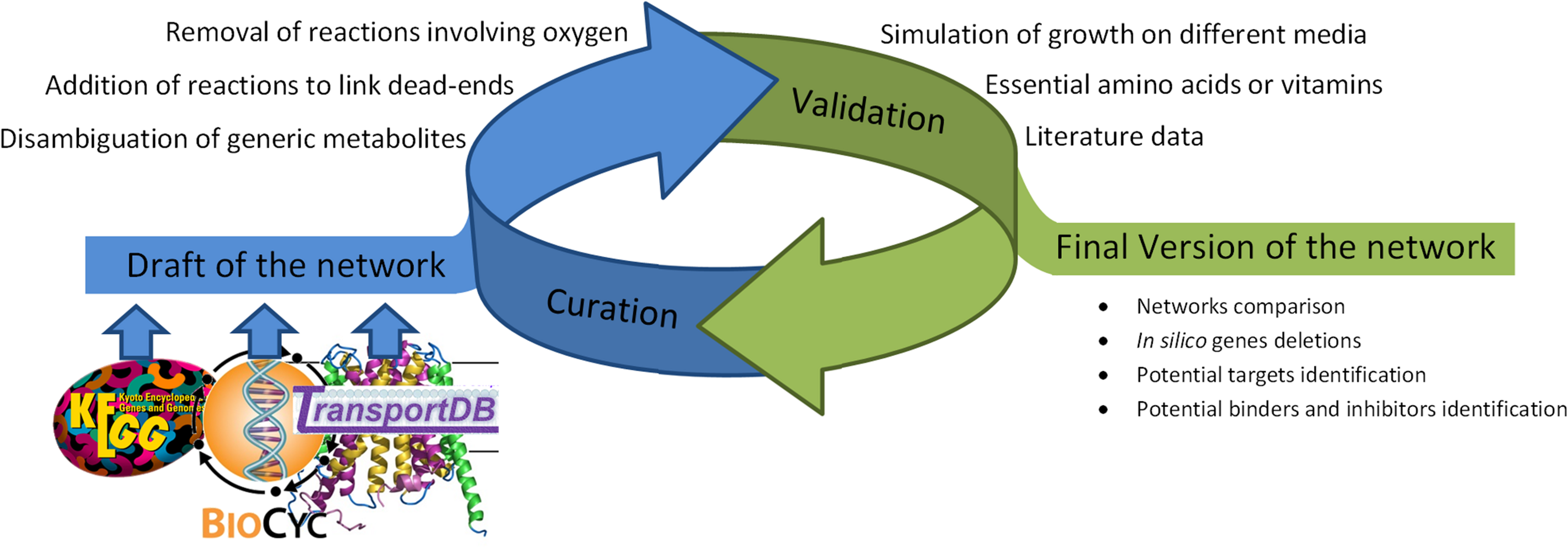
Flow chart representing the main steps involved in the metabolic network reconstruction. The creation of the network begins by the building of a draft by extracting data from various databases. The draft enters an iterative cycle involving curation and validation. Only growth simulations in different media and the effect of the removal of essential metabolites and vitamins as described in the literature are used to validate the network. The final version of the network was used to perform the various analyses. In particular, the comparison of predicted essential genes with the effect of their homologs in *B. subtilis* serves as further independent validation of the network.

Most transport reactions were based on TransportDB (Ren *et al*, 2007) although putative transporters not present in the databases had to be added for molecules known to be exported or imported by the bacteria. An exchange reaction (reaction that simulates interaction with the media via the appearance or disappearance of the given metabolites in the network) was created for each metabolite with an extracellular version. These exchange reactions are set to allow the export of a metabolite unless it is part of the tested growth medium in which case import is also possible.

The curation not only involved the addition and suppression of reactions from the initial draft, many characteristics of each reaction such as the directionality, the presence of complexes or the assignment of gene-reaction associations and their inclusion as part of a pathway had to be analyzed manually. The directionality of reactions was based on information obtained from the MetaCyc database when available. Reactions found exclusively in KEGG were kept bidirectional unless leading to the production of highly energetic compounds (ATP, NAD+, NADP+, etc.) that are not known to be produced in such a manner. Examples of reactions producing highly energetic compounds are ATP synthases, amino acid fermentation and glycolysis.

Protein-protein interactions are highly important in the network since they can modify the essentiality of a gene based on knowledge of the involvement of its protein product as part of a protein complex. The possibility that a protein complex is responsible for catalysis was investigated for every reaction that had more than one gene associated to the reaction and for any gene whose protein is identified as a subunit of a complex. We used information from TransportDB, UniProt, Brenda and literature data (either for the *C. difficile* protein of interests or a homolog of same function that could indicate a similar interaction between genes products). In every case the STRING version 9.1 (Franceschini *et al*, 2013) score was calculated to evaluate the confidence score attributed to each complex (Table E11).

We used KEGG pathway identifiers since they are generic and allow maintaining a small number of pathways in the network. Pathways containing less than five reactions were manually merged to the closest relevant pathway. Reactions that were not associated to any pathway in KEGG (32% of the reactions in KEGG) were linked to existing pathways. The pathway assigned to the reaction was the one in which the less linked metabolite of the reaction was most often present in the network (Table E9).

### Lipid biosynthesis and cell membrane composition

A problem with lipid biosynthesis in metabolic networks is the fact that a variety of different fatty acids exist in cells with different lengths and saturation states and can be used in numerous cellular processes, both metabolic and non-metabolic. Also, the lipid composition of the cellular membrane is a mix of various phospholipids and glycerolipids of varying lengths and composition and this composition varies depending on the growth conditions (Evans *et al*, 1998). A complete definition of lipid metabolism is likely impossible to define at this moment given the lack of experimental information specific to *C. difficile* membranes. Furthermore, the resulting network would be dominated by lipid reactions with the same few genes repeated for every possible length and saturation state for every lipid type. Therefore, having an exhaustive definition of lipid metabolism would not bring any additional relevant information on metabolism. In most reconstructed metabolic networks, lipids are used almost exclusively in membrane formation. Following (Senger & Papoutsakis, 2008), in order to reduce the complexity of lipid metabolism while keeping it as close to the real bacteria as possible, the fatty acid composition of all lipids is held at a constant 16:0 (carbon chain length: number of double bonds), which is the most abundant fatty acid observed in *C. acetobutylicum* (Johnston & Goldfine, 1992). The last step in simplifying lipid biosynthesis was to combine the elongation of fatty acids from acetyl (2:0) to palmitate (16:0) into a single reaction and beta-oxidation of the palmitate back into acetyl as another single reaction.

### Biomass

In order to simulate bacterial growth, the biomass (ensemble of macromolecules necessary for cellular growth and division) composed of DNA, RNA, cell wall, proteins, solute pool and lipids was defined based on the following elements. Nucleic acid composition of DNA used in the network was calculated based on the nucleotide content of the genome and plasmid of *Clostridium difficile* strain 630, RNA composition is based on the content of the transcriptome using the UCSD genome browser (Chan *et al*, 2012) and protein composition using the proteome (Table E12). Lipid, cell wall and solute pool composition as well as the overall biomass composition were taken from the *C. acetobutylicum* network (Senger & Papoutsakis, 2008) due to a lack of literature specific to *C. difficile*.

### Simulation of growth and gene essentiality

Two methods were used to simulate bacterial growth and determine gene essentiality: flux balance analysis (FBA) (Orth *et al*, 2010) and synthetic availability (SA) (Wunderlich & Mirny, 2006).

#### Flux balance analysis (FBA)

Flux balance analysis (Orth *et al*, 2010) is a constraint-based modeling method commonly used in the study of genome-scale metabolic networks. FBA firsts creates a stoichiometry matrix (*S*) from the network where each row represents a metabolite and each column a reaction. The values in this matrix correspond to the stoichiometry of the metabolite in the reaction with a negative number representing consumption and a positive number representing production of the metabolite. A system of linear equations is produced by multiplying *S* with a column vector *v* representing the fluxes through each reaction. FBA creates a steady-state distribution of fluxes where the product of the previous multiplication must equal zero *S* · *v* = 0. Since the resulting system of linear equations is undetermined, FBA uses linear programming to maximize a particular objective function *Z*, in our case biomass as the representation of growth, by maximizing the multiplication of a row vector *c* containing the weight of each reaction on *Z* with the column vector *v* used previously (maximize *Z* = *c* · *v*). Values in *v* are constrained by lower and upper bounds representing various factors like enzyme directionality of the reaction, capacity, uptake, secretion rates, etc. FBA finds a distribution of fluxes in the network respecting the constraints on *v* and maximizing *Z* at the same time. In our studies, the Sybil package version 1.1.11 (Gelius-Dietrich *et al*, 2013) available for the free R environment for statistical computing (version 2.15.2) was used in order to run FBA simulations. Other tools also exist that can use FBA with different interfaces like the COBRA package (Schellenberger *et al*, 2011) that runs on the proprietary Matlab computing environment.

#### Synthetic accessibility (SA)

Synthetic accessibility (Wunderlich & Mirny, 2006) is a parameter-free method to predict the essentiality of genes through their deletions in metabolic networks. SA uses network topology to calculate the number of reaction steps needed to produce the outputs (biomass) of the network from the inputs metabolites available in the growth medium. SA works by examining all reactions that use only input metabolites and marks those reactions and their products as ‘accessible’. In an iterative manner, successive iterations search for new reactions that have all required substrates marked as accessible until no new reaction can be added. Each metabolite *j* have a synthetic accessibility value *S_j_* representing the number of iterations before this metabolite became accessible and the Synthetic Accessibility of the network as a whole, *S_net_*, is the sum of *S_j_* of each of the output metabolites. Increases in *S_net_* resulting from gene deletions are predicted as being deleterious. Synthetic accessibility is a simpler method than FBA and gives comparable results on the prediction of essential genes demonstrating that the topology of the network is the principal factor influencing essentiality. We use our own implementation of the algorithm.

## ACKNOWLEDGEMENTS

ML was the recipient of two undergraduate research fellowships from the Faculty of Medicine, Université de Sherbrooke. TC was partially funded by PROTEO (the Québec network for research on protein function, structure and engineering), the Fonds de Recherche du Québec en Santé (FRQ-S) and currently holds a CREMUS graduate fellowship from the Faculty of Medicine, Université de Sherbrooke. R.J.N. was a FRQ-S Junior 1 research fellowship holder and a member of the Centre de Recherche Clinique Étienne-Le Bel, Institute of Pharmacology of Sherbrooke, GRASP (Groupe de Recherche Axé sur la Structure des Protéines) and PROTEO.

## AUTHOR CONTRIBUTIONS

ML and TC created the reconstructed metabolic network and performed the single and double mutant experiments. ML performed the detection of homologs and their inhibitors. RN designed the experiments. ML, TC and RN wrote the manuscript.

## EXPANDED VIEW

**Figure E1: Comparison of the metabolites (A) and genes (B) in *i*MLTC804cdf and the automatically generated *C. difficile* network.** A vast number of unique metabolites in the automatic network (485 out of 487) are generic and do not contribute to a functional network. In *i*MLTC804cdf only 23 generic metabolites exist. Four metabolites in *i*MLTC804cdf are associated to multiple entries in the automatic network while 11 others do not even appear in *i*MLTC804cdf. Likewise, 382 loci are repeated in the automatic network, of which 304 appear as unique loci in *i*MLTC804cdf. Of 68 loci associated to non-metabolic functions, only 3 appear in *i*MLTC804cdf. A total of 21 predicted essential genes (6 as single mutant and 15 as a pair, small red circle) are absent in the automatic network. Numbers in parenthesis represent totals in each section of the Venn diagram and those in brackets separated by a forward slash represent *i*MLTC804cdf and the *C. difficile* automatically generated network respectively.

**Figure E2: Comparison of the metabolites (A), genes (B) and reaction (C) from the *Clostridium difficile* metabolic network *i*MLTC804cdf to those from the network of *Clostridium acetobutylicum*.** Numbers in parenthesis represent totals in each section of the Venn diagram and those in brackets separated by a forward slash represent *i*MLTC804cdf and the *C. acetobutylicum* network respectively. *i*MLTC804cdf has 65% more reactions, 29% more unique metabolites and 69% more unique loci than the curated *C. acetobutylicum* network.

**Figure E3: Examples of curation of reactions involving oxygen.** Some reactions (A) were simply removed from the network; others (B) were modified to use different electron acceptors after proof of the feasibility for this alternative version has been found in the literature while, in other cases, (C) full pathway of aerobic biosynthesis or degradation had to be replaced by their anaerobic version such as in the case of Vitamin B12 degradation. For panels B and C, reactions highlighted with red squares were the ones initially linked with the EC number while the reaction in the green squares is the modified version based on the Enzyme Commission annotation of the reaction for anaerobic species.

**Figure E4: Stickland reactions.** Stickland reactions are an example of reactions that are often incompletely annotated in metabolic databases. Stickland reactions were described in the literature for *Clostridium difficile* and thus added to the network. In these reactions, two Stickland donors (proline, glycine, hydroxyproline or in some case leucine) reduced two acceptors (denoted as A in bold) that can then be used by one Stickland acceptor (valine, serine, methionine, leucine, isoleucine, threonine, alanine, aspartate and phenylalanine) to keep the electron balance of the cell. Each fermented amino acid leads to the production of one molecule of ATP for a total of 3 if all reactions are used (2 donors, 1 acceptor).

The uploaded zip file contains all extended materials as follows:

- Figure E1 (TIFF format)
- Figure E2 (TIFF format)
- Figure E3 (TIFF format)
- Figure E4 (TIFF format)
- Table E1 (Excel format)
- Table E2 (Excel format)
- Table E3 (Word format)
- Table E4 (Word format)
- Table E5 (Excel format)
- Table E6 (Excel format)
- Table E7 (Excel format)
- Table E8 (Excel format)
- Table E9 (Excel format)
- Table E10 (Excel format)
- Table E11 (Excel format)
- Table E12 (Excel format)
- Table E13 (Excel format)
- iMLTC804cdf.tsv (TSV formatted metabolic network)
- iMLTC804cdf.xlxs (Excel formatted metabolic network)
- iMLTC804cdf.xml (SBML v. 2.0 formatted metabolic network)

Please refer to the uploaded zip file to view this content.

